# Piezo activity levels need to be tightly regulated to maintain normal morphology and function in pericardial nephrocytes

**DOI:** 10.1101/2021.10.23.465463

**Authors:** Kristina Schulz, Paris Hazelton-Cavill, Karl K. Alornyo, Ilka Edenhofer, Maja Lindenmeyer, Christian Lohr, Tobias B. Huber, Barry Denholm, Sybille Koehler

## Abstract

Due to their position on glomerular capillaries, podocytes are continuously counteracting biomechanical filtration forces. Most therapeutic interventions known to generally slow or prevent the progression of chronic kidney disease appear to lower these biomechanical forces on podocytes, highlighting the critical need to better understand podocyte mechano-signalling pathways. Here we investigated whether the mechanotransducer Piezo is involved in a mechanosensation pathway in *Drosophila* nephrocytes, the podocyte homologue in the fly.

Loss of function analysis in Piezo depleted nephrocytes reveal a severe morphological and functional phenotype. Further, pharmacological activation of endogenous Piezo with Yoda1 causes a significant increase of intracellular Ca^++^ upon exposure to a mechanical stimulus in nephrocytes, as well as filtration disturbances. Elevated Piezo expression levels also result in a severe nephrocyte phenotype. Interestingly, expression of Piezo which lacks mechanosensitive channel activity, does not result in a severe nephrocyte phenotype, suggesting the observed changes in Piezo wildtype overexpressing cells are caused by the mechanosensitive channel activity. Moreover, blocking Piezo activity using the tarantula toxin GsMTx4 reverses the phenotypes observed in nephrocytes overexpressing Piezo.

Taken together, here we provide evidence that Piezo activity levels need to be tightly regulated to maintain normal pericardial nephrocyte morphology and function.

## Introduction

Kidney podocytes are highly specialized epithelial cells, which together with the fenestrated endothelium and the glomerular basement membrane, constitute the three-layered, size and charge selective filter of the glomerular filtration barrier ^1^. Podocytes have a unique morphology, forming primary and secondary foot processes which completely enwrap the capillaries. Injury to podocytes causes severe morphological change, detachment from the basement membrane and finally loss of podocytes into the primary urine with pathological consequences as these cells are post-mitotic and cannot be replenished. Podocyte injury can occur because of a variety of mechanisms, among them genetic deficiencies ^2–6^, toxin-mediated triggers ^7^ and changes in the physical environment within the glomerulus ^8–11^.

Podocytes are under constant mechanical force and regularly face changes in these forces, as they are responsible for the filtration of ∼180 litres of primary urine each day. The main forces arising from this are hydrostatic pressure (50 kPa) and fluid shear stress at the filtration slits (8 Pa) ^12^. These forces increase during diseases such as diabetes and hypertension and result in podocyte loss ^10^. It has previously been shown, that increased biomechanical force such as glomerular hypertension and fluid shear stress can cause podocyte depletion ^13–15^. Further, it is known, that remaining podocytes respond by growing in size to re-cover the blank capillary to overcome the challenge originating from loss of neighbouring cells; this is considered to be an adaptive and protective mechanism ^14,16,17^. In order for this to happen, one possible hypothesis is that podocytes exhibit mechanotransductive mechanisms to recognize these changes in the physical environment and initiate adaptive and protective responses, likely candidates are TRPC6, YAP, Talin, Filamins and Piezo ^5,8,9,18–21^.

Piezo is a mechanosensitive ion channel that is involved in mechanotransduction in many developmental and physiological contexts. Piezo channels (Piezo1 and Piezo2) open in response to mechanical force/stimuli and allow the influx of positively charged ions such as Ca^++^ and Na^+^ ^22,23^. The two proteins have different functional roles as Piezo1 was reported to be essential for sensing blood flow-associated shear stress and blood vessel development ^24–26^, while Piezo2 responds to touch and proprioception ^27^.

Whether Piezo plays a role in mechanotransduction in nephrocytes, the podocyte homologue in flies, is unknown. To address this question, we investigated the effect of Piezo depletion, activation and overexpression on nephrocyte morphology and function. Nephrocytes form foot processes, which are spanned by the nephrocyte diaphragm, a highly-specialized cell-cell contact, that is highly similar to the mammalian filtration barrier in the glomeruli ^28,29^. They function as filtration units in the fly and filter toxins and waste products from the haemolymph. Our data show that Piezo functions as a mechanotransducer in nephrocytes and reveal the need for careful regulation of its expression levels to prevent a nephrocyte phenotype.

## Results

### Piezo1 is expressed in mammalian podocytes

To investigate the functional role of Piezo we utilized *Drosophila* nephrocytes, the homologue cells to mammalian podocytes. To confirm Piezo expression in nephrocytes, we analysed publicly available single nucleus RNAsequencing data from the fly kidney atlas, which shows expression of Piezo in garland nephrocytes (36.6% of cells) and pericardial cells (18.3% of cells) ^30^. Of note, in *Drosophila* only one Piezo isoform is present, with equal similarity to human Piezo1 and Piezo2. As Piezo specific antibodies for Drosophila are elusive, we decided to confirm the presence of the channel by loss of function analysis and using a specific activator of the channel combined with assessment of downstream signaling pathways.

However, we examined Piezo1 protein expression in mammalian tissue. Proteomic analysis of isolated murine podocytes revealed expression of Piezo1 (12.25% enrichment when compared to non-podocyte glomerular cells) ^31^, strengthening support for our hypothesis that Piezo1 is a mechanotransducer in mammalian podocytes as well. Moreover, recent studies confirmed expression of Piezo1 in podocytes and also showed increased Piezo1 expression in podocytes in experimental hypertensive nephropathy and lupus nephritis ^32,33^. We examined expression of Piezo1 in mammalian glomeruli, by co-staining with the podocyte marker Synaptopodin and found Piezo1 expression in podocytes **(Figure 1A)**. Of note Piezo1 seems to localize within foot processes as co-localisation with synaptopodin was observed, but also within the cell body. Investigating publicly available single nucleus RNAsequencing data for human kidney also revealed expression of Piezo1 in podocytes, which seems to decrease in diabetic kidney disease **(Figure 1B)** ^34^. Moreover, searching Piezo1 in the single cell RNAsequencing data from the Kidney Precision Medicine Project revealed a disease-associated expression of the channel (mean expression: healthy controls: 1.45; chronic kidney disease (CKD): 1.73).

**Figure 1:**
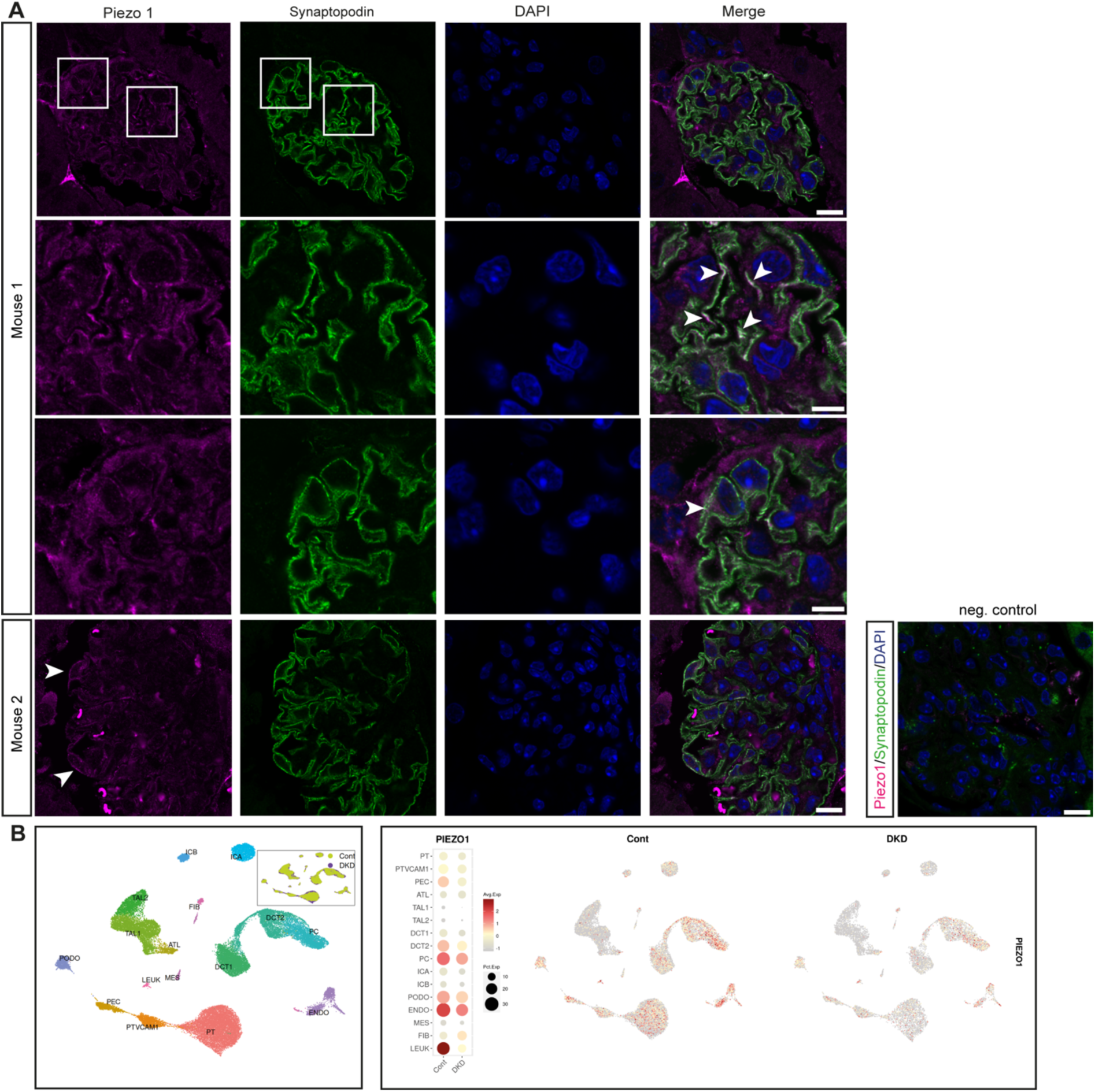
Expression of Piezo in mammalian tissue. **A** Piezo1 is expressed in mammalian podocytes, which are visualized using the podocyte marker Synaptopodin. Piezo1 is expressed in areas close to the slit diaphragm and in the cell body (arrow heads). Scale bar = 10μm. Negative control: without primary antibodies. Scale bar = 10μm. **B** Single nucleus RNAsequencing data from human diabetic kidney disease samples (DKD) and controls reveal Piezo1 expression in podocytes, which decreases in DKD. Data used from the Humphries lab with permission ^34^. PT: proximal tubule, PTVCAM1: VCAM1(+) proximal tubule, PEC: parietal epithelial cells, ATL: ascending thin limb, TAL1: CLDN16(-) thick ascending limb, Tal2: CLDN16(+) thick ascending limb, DCT1: early distal convoluted tubule, DT2: late distal convoluted tubule, PC: principal cells, ICA: type A intercalated cells, ICB: type B intercalated cells, PODO: podocytes, ENDO: endothelial cells, MES: mesangial cells and vascular smooth muscle cells, FIB: fibroblasts, LEUK: leukocytes.

Taken together, these data show Piezo1 is expressed in murine and human podocytes, localizes within the foot processes and the cell body and seems to have a disease-associated expression pattern.

### Loss of Piezo results in a morphological and functional nephrocyte phenotype

Due to confirmed expression of Piezo1 in podocytes by us and others ^21,32,33^, we then examined its role in nephrocytes. Within this study, we mainly focused on assessment of adult pericardial nephrocytes as this cell type is attached to the heart tube, which exhibits peristaltic movements and induces haemolymph movement. Thus, one can envision that adult pericardial nephrocytes experience some kind of biomechanical force *in vivo*. To unravel whether the mechanotransducer Piezo is present in nephrocytes, and moreover, is important for nephrocyte function, we used Piezo-specific RNAi for nephrocyte-specific depletion of Piezo. Here, two temperatures have been assessed, with increasing temperature causing increased activity of the nephrocyte-specific Gal4 driver, thus higher expression of Piezo-RNAi and increased knockdown efficiency. We investigated morphology of pericardial nephrocytes by visualizing the nephrocyte diaphragm using a Duf (dNEPH) and Pyd (dZO1) antibody, which reveals a finger-print like pattern in healthy nephrocytes. This resulted in a morphological phenotype, as nephrocyte diaphragm length is significantly reduced and the finger-print like pattern is less dense when compared to controls **(Figure 2A)**. Expression of Piezo-RNAi at 28°C, which increases knockdown efficiency, also results in a significant shortening of the nephrocyte diaphragm, which is more severe in comparison to 25°C **(Figure 2B)**. In addition to nephrocyte morphology, we also assessed nephrocyte function by performing FITC-Albumin uptake assays. FITC-Albumin can be filtered through the nephrocyte diaphragm and taken up by nephrocytes and accumulates in the lacunae prior to endocytosis. Changes in nephrocyte diaphragm integrity, subsequent alterations in the lacunae channels and endocytosis therefore result in altered levels of FITC-Albumin in the lacunae system. In line with the morphological phenotype, we also observed a severe and significant reduction of FITC-Albumin in Piezo depleted nephrocytes at 25°C and 28°C, reflecting a functional defect in adult pericardial cells **(Figure 2C,D)**.

**Figure 2:**
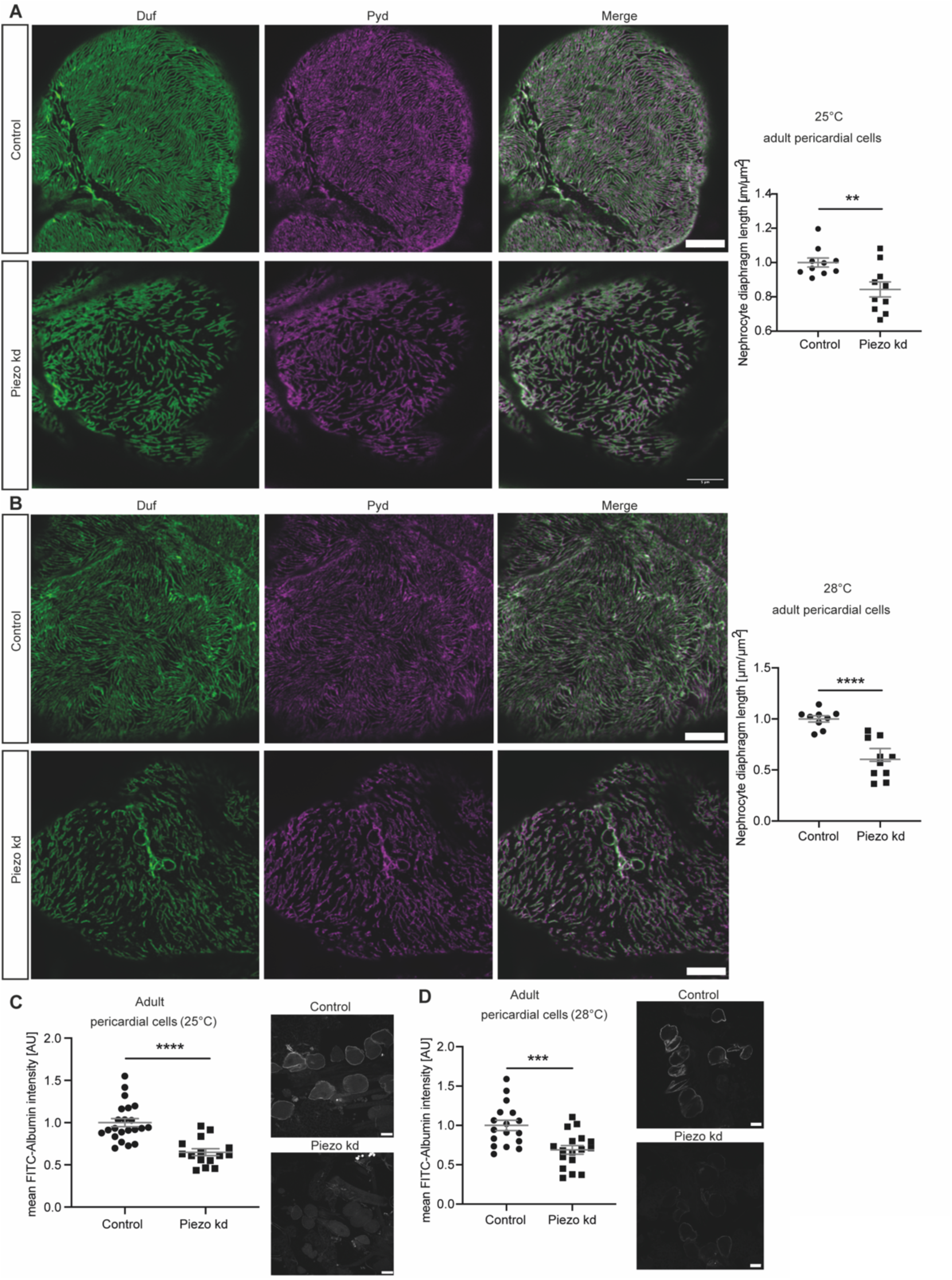
Loss of Piezo causes a severe nephrocyte phenotype in adult pericardial cells. **A** Visualization of the nephrocyte diaphragm with Duf and Pyd antibodies reveals a significant decrease in nephrocyte diaphragm length in adult pericardial cells upon loss of Piezo at 25°C (Piezo kd). Student’s t-test: **: p<0.01. Scale bar = 5μm. Control: w;*sns*-Gal4/+;+/UAS-*dicer2*; Piezo kd: w;*sns*-Gal4/+;UAS-*piezo*-RNAi/UAS-*dicer2*. **B** Immunofluorescence staining with a Duf and Pyd antibody reveal significant morphological changes in Piezo kd adult pericardial cells in comparison to control cells at 28°C. Scale bar = 5μm. Student’s t-test: ****: p<0.0001. **C** FITC-Albumin uptake assays in adult pericardial cells show a significant reduction of FITC-Albumin uptake in Piezo kd cells at 25°C. Scale bar = 25μm. Student’s t-test: ****: p<0.0001. **D** At 28°C FITC-Albumin uptake assays show a significant filtration defect in adult pericardial Piezo kd cells. Scale bar = 25μm. Student’s t-test: ***: p<0.001.

Taken together, our data shows that loss of Piezo results in a morphological and functional phenotype in adult pericardial cells. Further, the loss of function analysis also supports our hypothesis of Piezo expression in nephrocytes.

### Pharmacological activation of Piezo results in an increased Ca^++^ influx and filtration disturbances after applying a mechanical stimulus

Next, we aimed to understand Piezo’s role at physiological levels and when exposed to mechanical stress. Hence, we pharmacologically manipulated Piezo by using Yoda1, which specifically facilitates Piezo opening by reducing its mechanical activation threshold ^35,36^. The activation of the endogenous channel with the specific activator Yoda1 will further confirm its presence in nephrocytes.

As Piezo is a Ca^++^ permeable cation channel and sensitive to mechanical stimuli and stretch, we investigated the effect of Yoda1 by measuring Ca^++^ levels in adult pericardial cells after applying a mechanical stimulus. In detail, wildtype nephrocytes were incubated with either Yoda1 or the equivalent volume of DMSO and exposed to a mechanical stimulus by applying a pressure-controlled puff of the corresponding solution (Yoda1 or DMSO, 0.2bar for 3secs). The applied pressure stimulus (puff) elicited a Ca^++^ signal in adult pericardial cells, which was significantly higher after incubation in Yoda1 when compared to control conditions. This suggests that Yoda1 enhances the mechanosensitive channel activity of endogenous Piezo and amplifies calcium signalling in these cells after applying a mechanical stimulus **(Figure 3A)**.

**Figure 3:**
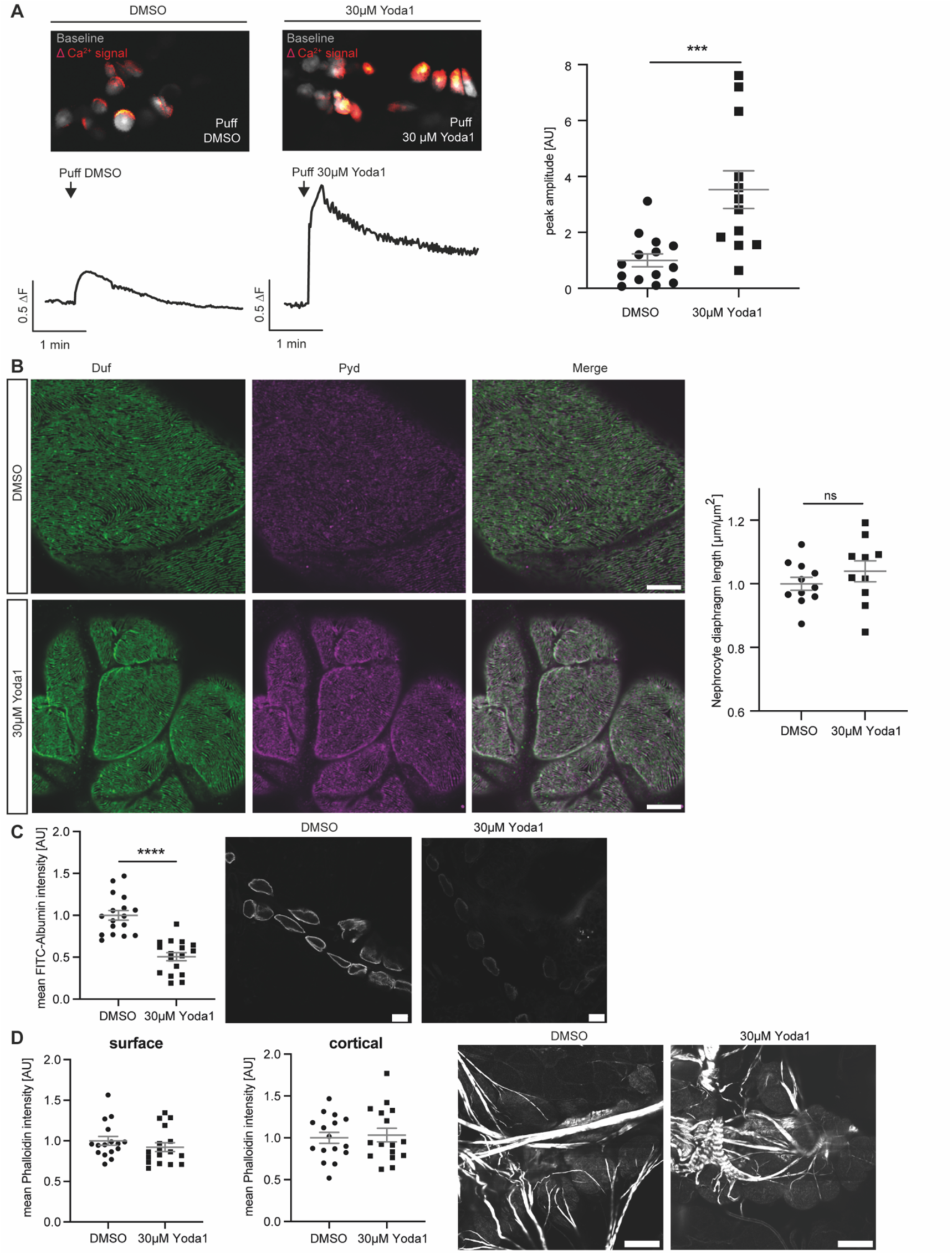
Pharmacological activation of Piezo with Yoda1 in adult pericardial nephrocytes. **A** Images of GCaMP6-fluorescent recordings showing puff-induced calcium signals (red) compared to baseline fluorescence (grey) in adult pericardial cells treated with DMSO (left) or 30µM Yoda1 (right). Below the images are corresponding traces of calcium signal changes over time, following a puff of DMSO (left trace) or 30µM Yoda1 (right trace). For analysis of GCamP6 intensity, ROIs of single cells have been analysed over time. Fluorescence intensity was then calculated as the mean for each fly and the maximum value within 25secs after puff is depicted as the peak amplitude. Yoda1 treated cells present with a significant increase in calcium signal amplitude after applying force (puff of Yoda1) when compared to DMSO-treated control cells (right). DMSO: 14 flies, 95 ROIs; Yoda1: 12 flies, 101 ROIs. Student’s t-test: ***: p < 0.001. w; *sns*-Gal4/+;UAS-*dicer2*/UAS-*GCaMP6m*. **B** 5mins incubation with Yoda1 in combination with applying a mechanical stimulus does not reveal any morphological changes as depicted by immunofluorescence staining with Duf and Pyd antibodies in adult pericardial cells. Scale bar = 5μm. **C** FITC-Albumin uptake assays reveal a significant difference between controls (DMSO) and Yoda1 treated cells, which have been exposed to a mechanical stimulus (puff). Student’s t-test: ****: p < 0.0001. Scale bar = 25μm. **D** Visualisation of the actin cytoskeleton with phalloidin does not reveal any changes upon Yoda1 treatment and mechanical stimuli. Scale bar = 25μm.

It is known that nephrocytes also elicit spontaneous calcium signals (SCSs) ^37^. We observed similar SCSs in only 5.88% of pericardial cells, whereas 87.65% of cells showed precisely timed and synchronous puff-induced calcium signals **(Supp. Figure 1A,B)**. The puff-induced signals differed significantly from SCSs in shape and timing, which was confirmed by quantification of both **(Supp. Figure 1C,D,E)**.

Moreover, Yoda1 treatment in combination with the mechanical stimulus revealed a functional defect in wildtype adult pericardial cells with significantly decreased FITC-Albumin accumulation **(Figure 3C)**, while morphology was unaffected **(Figure 3B)**.

Ca^++^ has an impact on actin fibre formation, hence we investigated the actin-cytoskeleton by using phalloidin, which binds to actin fibres and enables visualisation of those. Although nephrocytes show an increased Ca^++^ influx in the presence of Yoda1 and applying mechanical stimuli, no differences in cortical and surface actin fibre formation could be observed in comparison to control cells within the time frame of the experiment **(Figure 3D)**.

To confirm that the observed phenotypes are elicited by activation of endogenous Piezo in nephrocytes through Yoda1, we performed functional assessment of Piezo knockdown nephrocytes after Yoda1 treatment and application of the mechanical stimulus. We assessed Ca^++^ influx in these cells, revealing that while Yoda1 treated wildtype cells showed a significant increase of Ca^++^ signals after applying a mechanical stimulus, this increase was significantly less in Yoda1 treated Piezo knockdown cells and comparable to wildtype cells treated with DMSO **(Figure 4A)**.

**Figure 4:**
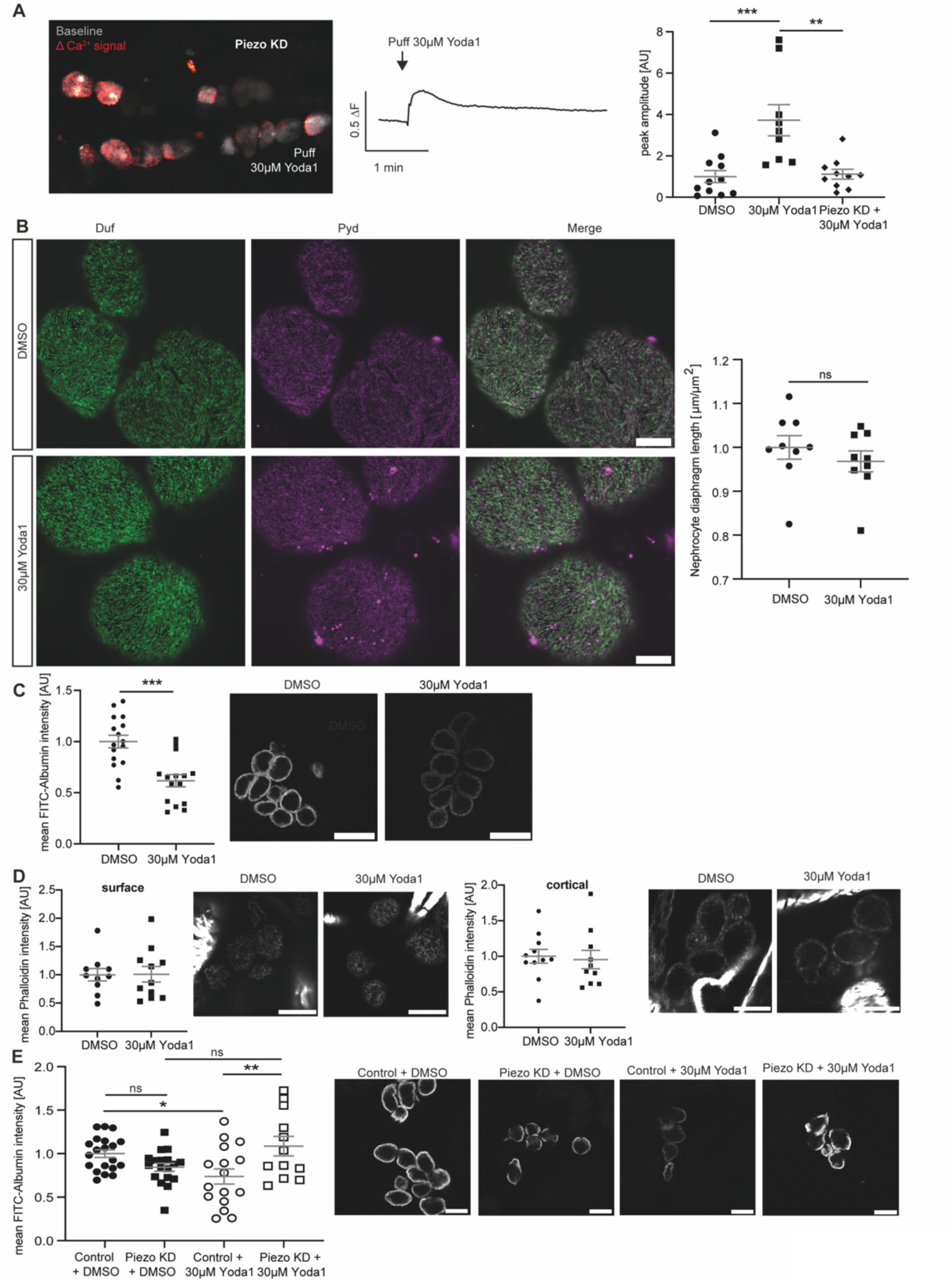
Pharmacological activation of Piezo with Yoda1 and applying mechanical force results in filtration disturbances, which are abrogated upon Piezo depletion in garland nephrocytes. **A** Calcium imaging reveals an abrogated calcium influx in Yoda1-treated Piezo kd pericardial cells. Images display GCamP6-fluorescent recordings of puff-induced calcium signals (red) compared to baseline fluorescence (white) in adult pericardial cells (left). The corresponding trace shows the calcium signal over time following a puff of 30µM Yoda1 (middle). Quantification is represented as peak amplitude (right) of puff-induced calcium signals in pericardial cells from wildtype flies treated with Yoda1 (30µM), DMSO (equivalent volume) and Piezo knockdown (KD) flies treated with Yoda1 (30µM). This reveals that puff-induced calcium signals in Piezo kd cells are comparable to those in DMSO-treated wildtype cells, while Yoda1-treated wildtype cells show a significant increase in calcium signals when subjected to mechanical stress. DMSO: 11 flies, 79 ROIs; Yoda1: 9 flies, 70 ROIs; Piezo KD: 10 flies, 69 ROIs. 1way-ANOVA with Tukey’s multiple comparison test: **: p < 0.01; ***: p < 0.001. **B** 5mins incubation with Yoda1 do not reveal any morphological changes as depicted by immunofluorescence staining with a Duf and Pyd antibody. Scale bar = 5μm. **C** FITC-Albumin uptake assays reveal a significant reduction of FITC-Albumin uptake after Yoda1 treatment and applying mechanical force. Scale bar = 25μm. Student’s t-test: ***: p<0.001. **C** Visualisation of the actin cytoskeleton with phalloidin does not reveal any changes upon Yoda1 treatment and applying mechanical force. Surface and cortical levels of actin fibres have been assessed. Scale bar = 15μm (phalloidin). **D** FITC-Albumin uptake assays reveal the absence of a phenotype after Yoda1 treatment and applying mechanical force in Piezo kd nephrocytes (w;*sns*-Gal4/+;UAS-*piezo*-RNAi/UAS-*dicer2*) when compared to control cells (w;*sns*-Gal4/+;+/UAS-*dicer2*). Scale bar = 25μm. 1way-ANOVA with Tukey’s multiple comparison test: *: p < 0.05; **: p < 0.01.

**Supp. Figure 1:**
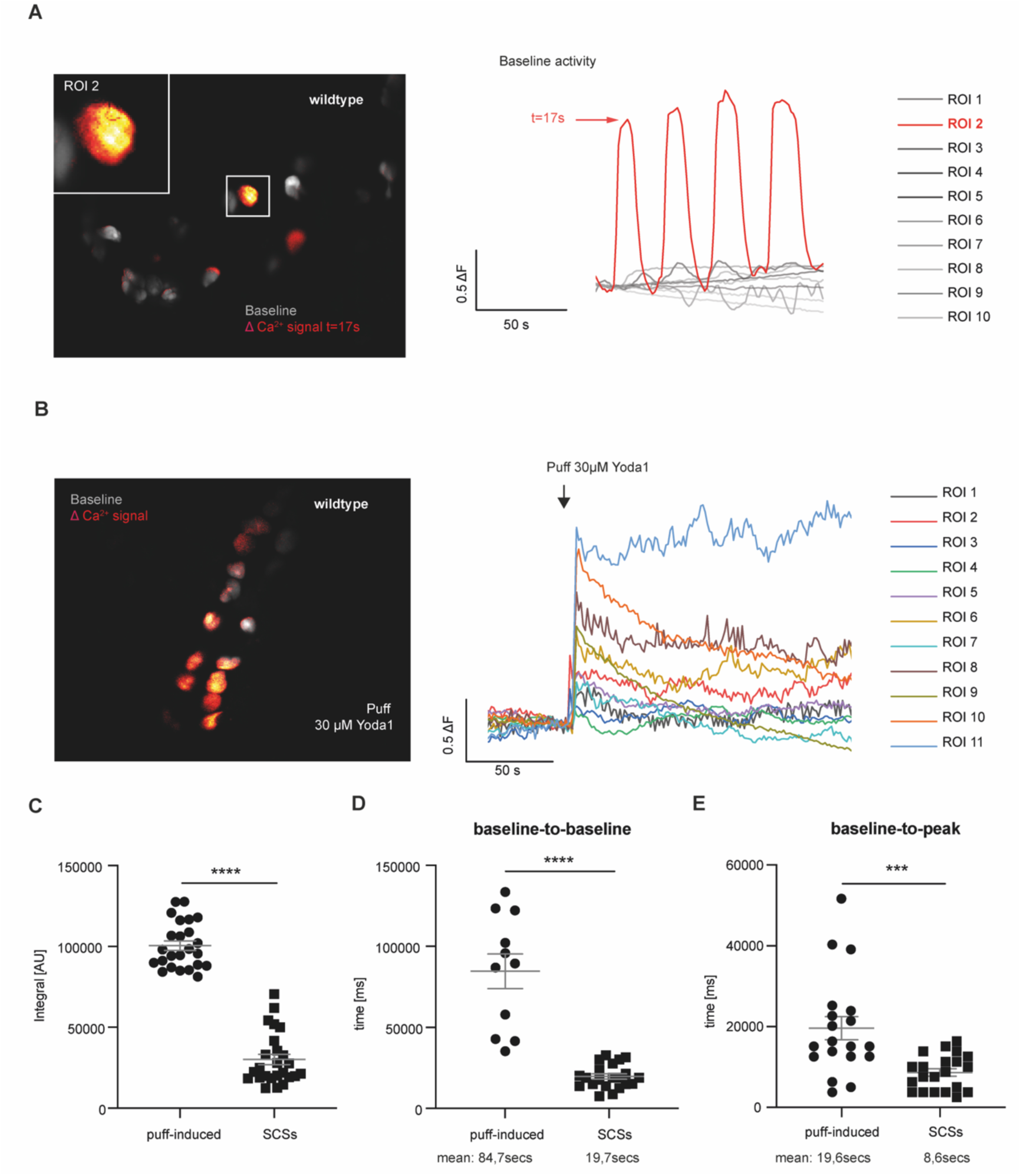
Calcium imaging in pericardial nephrocytes. **A** The image (left) shows spontaneous Ca^++^ activity (red) in GCamP6 expressing nephrocytes. Each pericardial cell was marked as an ROI and changes in calcium levels over the course of an experiment are depicted in corresponding traces (right). ROI2 shows the occurrence of spontaneous calcium signals. **B** The image (left) shows synchronous puff-induced calcium signals (red) in the majority of GCamP6-expressing nephrocytes and the corresponding traces of each ROI (right). **C** Quantification of the size of the integral of puff-induced (mean of all ROIs in one fly represent a datapoint, 23 flies in total) and spontaneous calcium signals (every SCSs in 11ROIs in 9 flies).; Student’s t-test: ****: p < 0.0001. **D** Quantification of baseline-to-baseline duration of puff-induced (mean of 11 flies, in which signals went back to baseline) and spontaneous calcium signals (every SCSs in 11ROIs in 9 flies). Student’s t-test: ****: p < 0.0001. **E** Quantification of baseline-to-peak duration in puff-induced (mean of 19 flies) and spontaneous calcium signals (every SCSs in 11ROIs in 9 flies). Student’s t-test: *** p < 0.001.

Since the knockdown of Piezo in adult pericardial nephrocytes results in a functional phenotype in the FITC-Albumin uptake assay itself, we could not assess the absence of this phenotype after applying Yoda1 and a mechanical stimulus to Piezo knockdown cells. Hence, we investigated Piezo loss in larval garland cells, which are similar to pericardial cells, but not identical. By doing so, we did not observe a functional phenotype **(Figure 4E)**. We also assessed the effects of Yoda1 treatment and applying a mechanical stimulus on wildtype larval garland nephrocytes and observed similar effects as in adult pericardial cells; a significant filtration defect, but no difference in morphology and the actin cytoskeleton **(Figure 4B,C,D)**. Treating Piezo knockdown garland nephrocytes with Yoda1 and applying a mechanical stimulus did not cause significant changes in FITC-Albumin uptake, when compared to DMSO treated Piezo knockdown cells or DMSO treated wildtype nephrocytes, suggesting that the decreased FITC-Albumin uptake in Yoda1 treated wildtype cells is mediated by Piezo **(Figure 4E)**.

Taken together, pharmacological activation of endogenous Piezo in nephrocytes causes an increased Ca^++^ influx and a functional defect after applying mechanical stimuli, while nephrocyte morphology and actin fibres were not altered within the timeframe of this experiment. The observed phenotypes upon Yoda1 treatment and applying a mechanical stimulus can be abrogated by depleting Piezo, confirming the presence and functional role of the channel in nephrocytes.

### Elevation of Piezo levels cause a nephrocyte phenotype, which is mediated by the mechanosensitive domain

Previously published data show an increase of Piezo1 levels in podocytes in a hypertensive nephropathy mouse model and lupus nephritis ^32,33^. Also, searching publicly available human data (KPMP) revealed an increase of Piezo1 in podocytes in chronic kidney disease. Hence, we further investigated whether an overexpression of the channel also results in a nephrocyte phenotype. To unravel the effect of the mechanosensitive domain of Piezo in nephrocytes in greater detail, we used two fly strains, either expressing Piezo wildtype (Piezo WT) or a Piezo mutant lacking the mechanosensitive channel activity (Piezo w/o MS) ^38^. Both variants localise to the cell cortex close to Pyd and within the lacunae system **(Supp. Figure 2)**. We observed a morphological phenotype with decreased nephrocyte diaphragm density in adult pericardial cells expressing Piezo wildtype at medium levels (25°C) **(Figure 5A; Supp. Figure 3A)**. Interestingly, expression of the mutant lacking the mechanosensitive channel activity did not cause any morphological changes at medium levels **(Figure 5A; Supp. Figure 3A)**. To assess whether a further increase of protein levels, in particular of the mutant, would cause a nephrocyte phenotype, we repeated morphological assessment at 28°C (high expression levels). While pericardial nephrocytes expressing Piezo wildtype exhibited a significant nephrocyte phenotype, expression of the mutant still did not cause any obvious changes at 28°C **(Figure 5B; Supp. Figure 3B)**.

**Figure 5:**
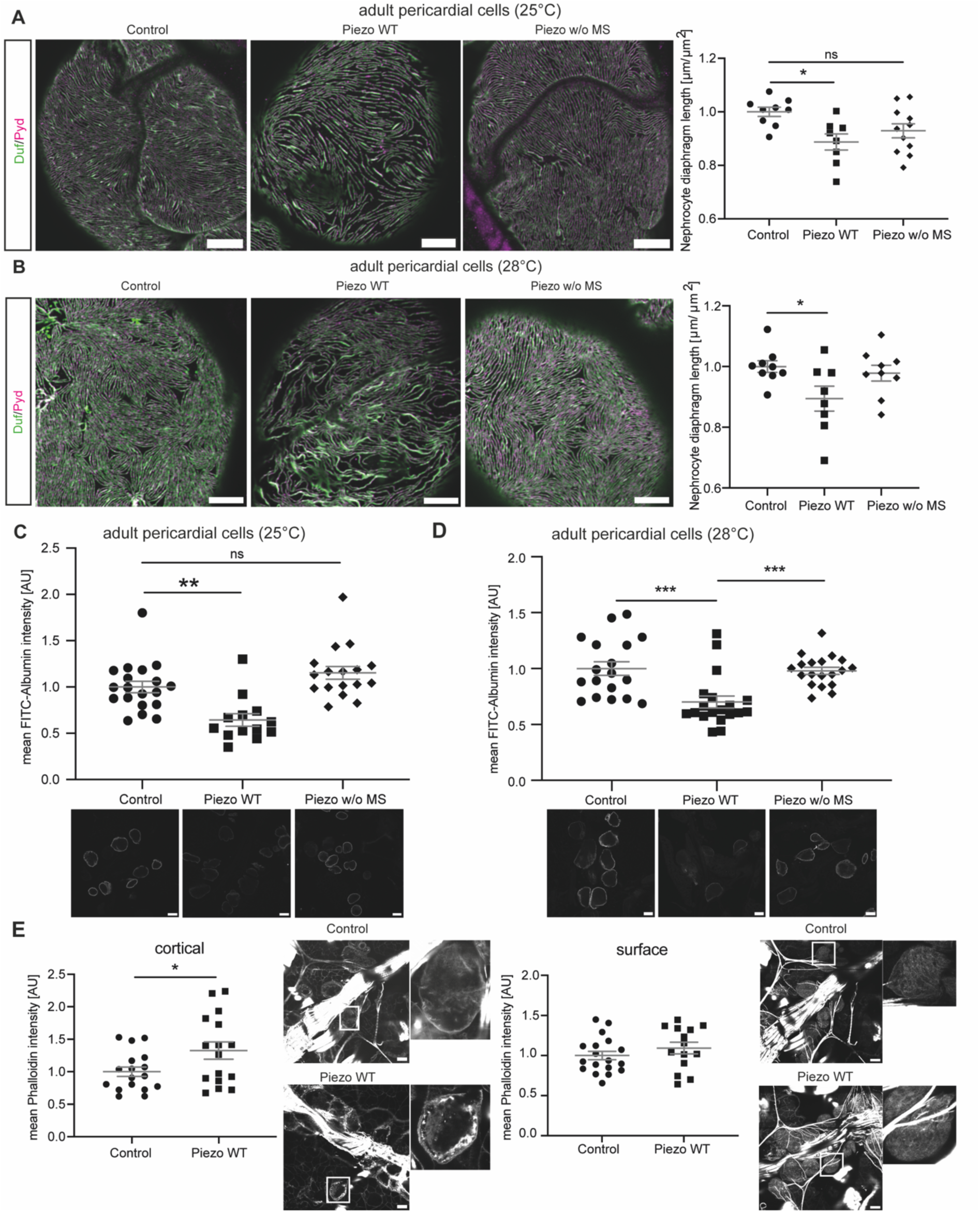
Elevated Piezo levels cause a severe nephrocyte phenotype in adult pericardial nephrocytes, which is mediated by the mechanosensitive domain. **A** Visualization of the nephrocyte diaphragm with Duf and Pyd antibodies shows a significant reduction of nephrocyte diaphragm length in adult pericardial cells expressing Piezo wildtype, while cells expressing the mutant without the mechanosensitive channel activity do not exhibit a phenotype. Experiments have been done at 25°C resulting in medium expression levels. Scale bar = 5μm. 1way-ANOVA with Tukey’s multiple comparison test: *: p < 0.05. Control: w; *sns*-Gal4/+;UAS-*dicer2*/+. Piezo WT: w; *sns*-Gal4/+;UAS-*dicer2*/UAS-*piezo*-flag. Piezo w/o MS: w; *sns*-Gal4/+;UAS-*GCaMP6m*/UAS-*piezo*-2306MYC.flag. **B** Immunofluorescence staining to visualize the nephrocyte diaphragm at 28°C (high expression levels) also reveal a severe and significant phenotype upon expression of Piezo wildtype, while pericardial cells expressing the mutant exhibit a normal morphology. Scale bar = 5μm. 1way-ANOVA with Tukey’s multiple comparison test: *: p < 0.05. **C,D** FITC-Albumin uptake assays reveal a severe phenotype in adult pericardial cells overexpressing wildtype Piezo at 25°C (medium levels) and 28°C (high levels). Expression of the mutant variant of Piezo does not reveal a functional defect, neither at 25°C nor at 28°C. Scale bar = 25μm. 1way-ANOVA with Tukey’s multiple comparison test: **: p < 0.01; ***: p < 0.001. **E** Visualization of actin fibres by phalloidin immunofluorescence reveals a significant increase of cortical actin fibres in Piezo wildtype overexpressing adult pericardial cells, while surface levels are not altered. Scale bar = 25μm. Student’s t-test: *: p < 0.05.

**Supp. Figure 2:**
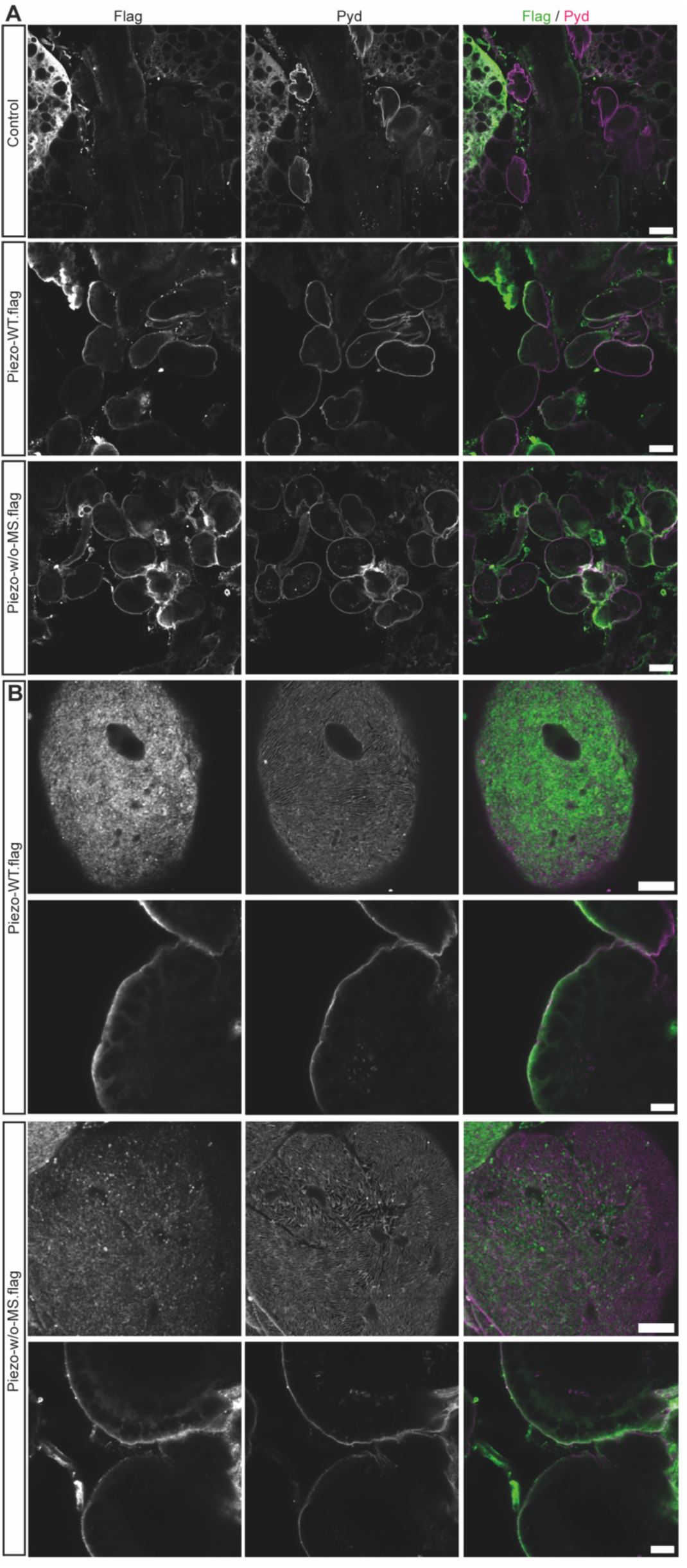
Overexpression of Piezo wildtype and a Piezo mutant lacking the mechanosensitive domain. **A** Immunofluoresence of adult pericardial nephrocytes expressing Piezo wildtype or the mutant Piezo at 25°C visualizing the nephrocyte diaphragm with a Pyd antibody and the flag tag to visualize Piezo. Scale bar = 25μm (upper three rows) and 5μm. Control: w; *sns*-Gal4/+;UAS-*dicer2*/+. Piezo WT: w; *sns*-Gal4/+;UAS-*dicer2*/UAS-*piezo*-flag. Piezo w/o MS: w; *sns*-Gal4/+;UAS-*GCaMP6m*/UAS-*piezo*-2306MYC.flag.

**Supp. Figure 3:**
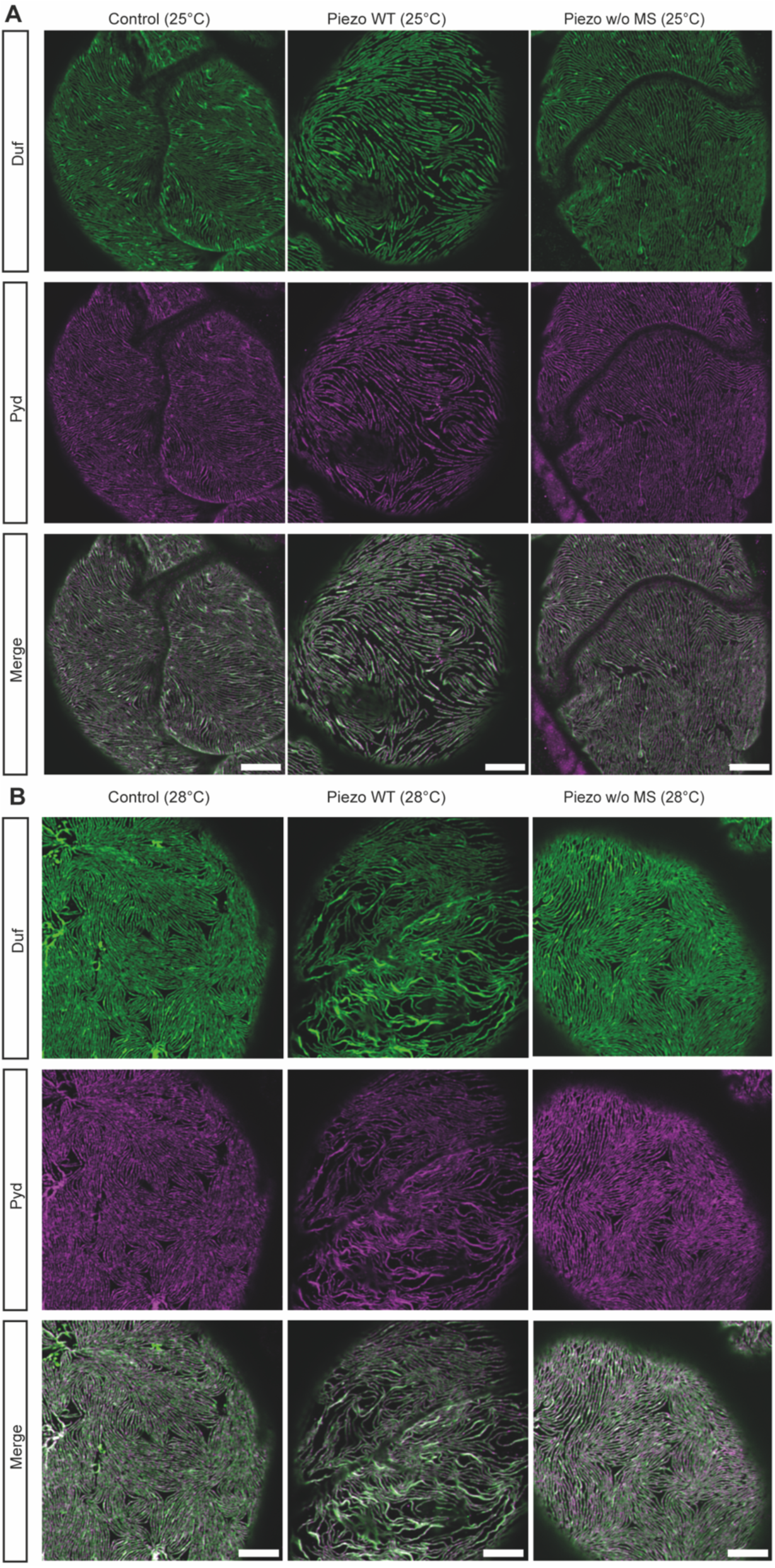
Overexpression of Piezo causes a nephrocyte phenotype, which is mediated by the mechanosensitive channel domain. **A,B** Immunofluoresence of adult pericardial nephrocytes expressing Piezo wildtype or the mutant Piezo at 25°C (medium levels) and 28°C (high levels) visualizing the nephrocyte diaphragm with Duf and Pyd antibodies. Scale bar = 5μm. Control: w; *sns*-Gal4/+;UAS-*dicer2*/+. Piezo WT: w; *sns*-Gal4/+;UAS-*dicer2*/UAS-*piezo*-flag. Piezo w/o MS: w; *sns*-Gal4/+;UAS-*GCaMP6m*/UAS-*piezo*-2306MYC.flag.

Further, we investigated nephrocyte function and observed significantly decreased FITC-Albumin uptake upon expression of Piezo wildtype at 25°C and 28°C, confirming a severe functional defect **(Figure 5C,D)**. We also assessed pericardial nephrocytes expressing medium and high levels of the Piezo mutant regarding FITC-Albumin uptake and did not observe any significant differences when compared to controls **(Figure 5C,D)**.

Although we did not observe alterations of the actin cytoskeleton by activating Piezo with Yoda1 (short-term effect), we investigated whether overexpression of Piezo wildtype (long-term effect) causes changes in actin fibre formation. Visualization by phalloidin and quantification of cortical actin fibres revealed a significant increase, while surface levels of actin fibres are unchanged **(Figure 5E)**.

Taken together, overexpression of Piezo is detrimental for nephrocyte biology and the observed morphological and functional phenotype seem to be mediated by its mechanosensitive domain.

### Pharmacological inhibition with tarantula toxin reverses the nephrocyte phenotypes observed upon overexpression of Piezo

Overexpression of Piezo has a detrimental effect on nephrocyte biology. Therefore, we tested whether pharmacological inhibition of the channel could reverse the phenotypes we observed. To do so, we used tarantula toxin (GsMTx4), which integrates into the cell membrane and blocks the opening of the channel upon mechanical stretch ^39^. Of note, GsMTx4 is not specific for inhibiting Piezo, but also acts on other cation channels.

First, we assessed whether treatment of pericardial cells with GsMTx4 results in a nephrocyte phenotype. Analysing morphology and function, we did not observe any significant changes in wildtype flies **(Figure 6A,B,C,D).** Next, we investigated whether a 5mins incubation of pericardial nephrocytes with the toxin reverses the morphological phenotype observed upon overexpression of Piezo wildtype. Adult pericardial cells present with a significant rescue of the morphological phenotype after GsMTx4 treatment **(Figure 6A,B; Supp. Figure 4)**. Moreover, FITC-Albumin uptake assays also reveal a significant rescue of the functional defect induced by Piezo overexpression after GsMTx4 treatment **(Figure 6C,D)**.

**Figure 6:**
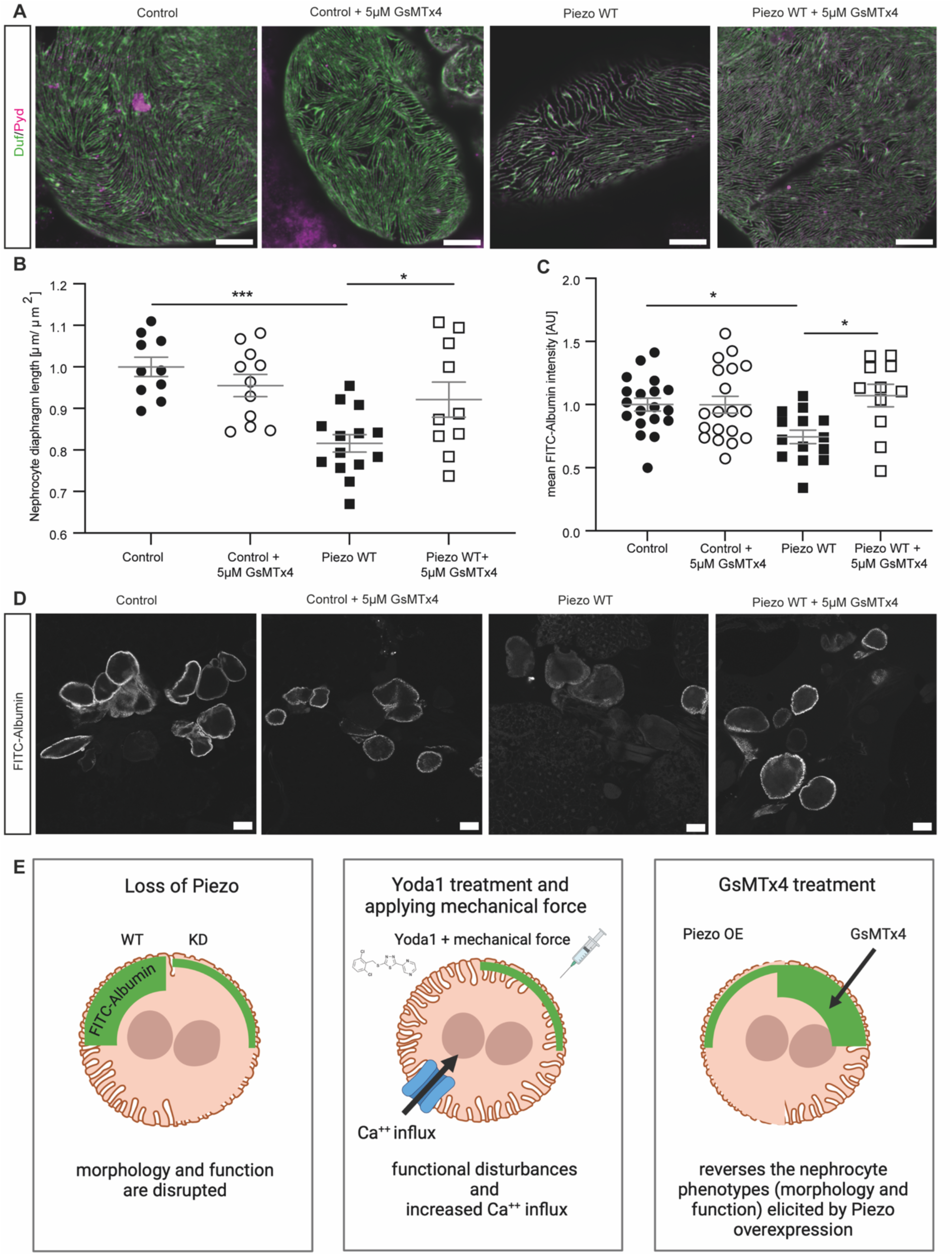
GsMTx4 treatment reverses the phenotypes observed in nephrocytes overexpressing Piezo wildtype. **A,B** Immunofluorescence staining with Duf and Pyd antibodies reveals a rescue of the morphological changes in adult pericardial cells expressing Piezo wildtype after incubation with 5μM GsMTx4 for 5mins. 1way-ANOVA with Tukey’s multiple comparison test: *: p < 0.05; ***: p < 0.001. Control: w; *sns*-Gal4/+;UAS-*dicer2*/+. Piezo WT: w; *sns*-Gal4/+;UAS-*dicer2*/UAS-*piezo*-flag. **C,D** FITC-Albumin uptake assays show a rescue of the observed phenotype in adult pericardial nephrocytes overexpressing Piezo wildtype after 5mins GsMTx4 treatment. 1way-ANOVA with Tukey’s multiple comparison test: *: p < 0.05. **E** Schematic summarising the data. First panel: Loss of Piezo causes a morphological and functional phenotype, which is most severe in adult pericardial cells at 28°C. Second panel: Pharmacological activation of endogenous Piezo by Yoda1 treatment and applying mechanical force causes functional defects and increased Ca^++^ influx. Third panel: Overexpression of wildtype Piezo in adult pericardial nephrocytes results in morphological and functional disturbances. These phenotypes seem to be mediated by the mechanosensitive channel activity of Piezo. Treatment of Piezo wildtype overexpressing nephrocytes with GsMTx4 reversed the observed phenotypes. All images have been created with BioRender.com.

**Supp. Figure 4:**
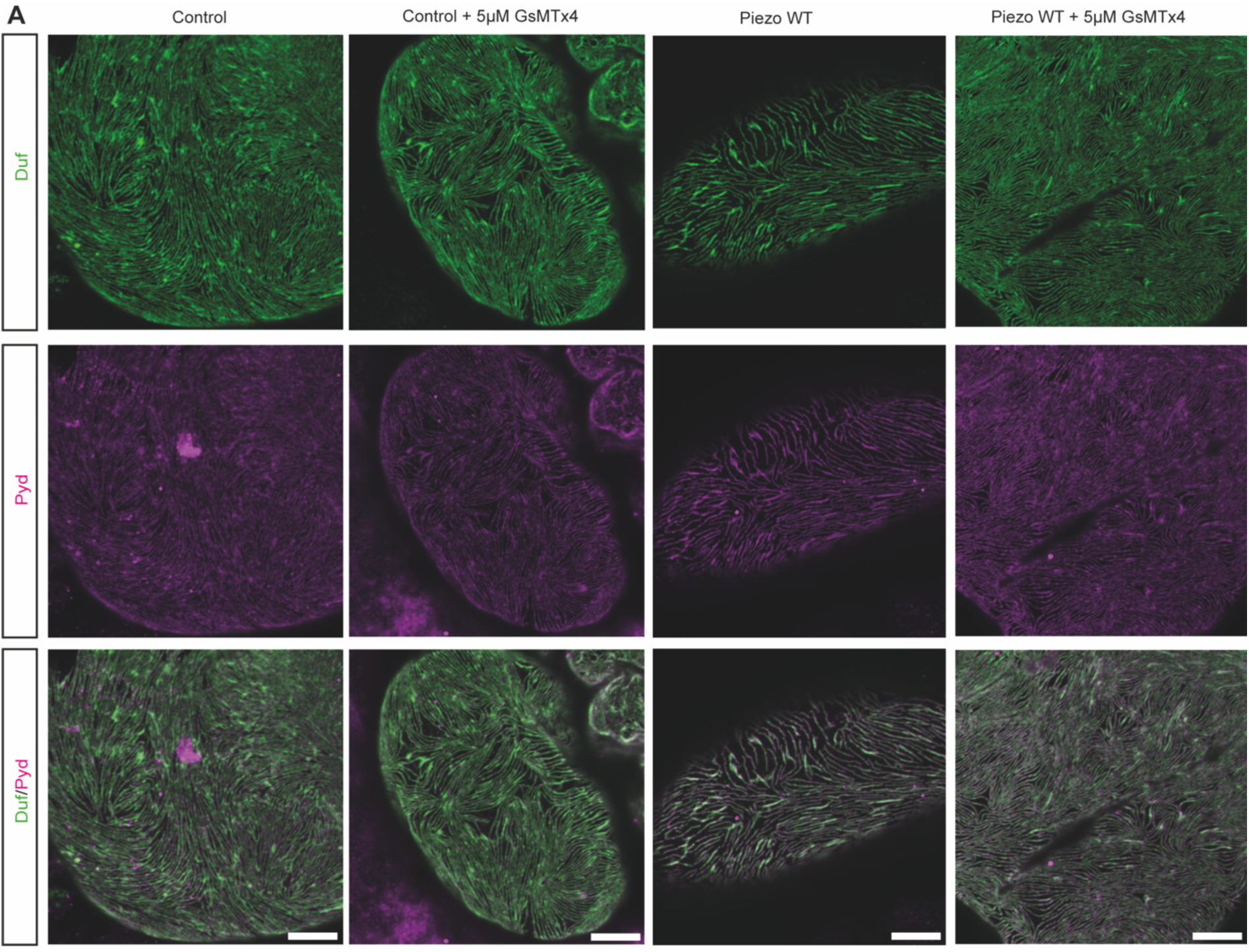
GsMTx4 treatment reverses the morphological phenotype observed in adult pericardial nephrocytes overexpressing Piezo wildtype. **A** Immunofluorescence staining with Duf and Pyd antibodies reveals a rescue of the morphological changes after incubation with 5μM GsMTx4 for 5mins in adult pericardial cells expressing Piezo wildtype. Scale bar = 5μm. Control: w; *sns*-Gal4/+;UAS-*dicer2*/+. Piezo WT: w; *sns*-Gal4/+;UAS-*dicer2*/UAS-*piezo*-flag.

Taken together, a treatment with the toxin did not reveal pathological effects on nephrocyte morphology and function, but reversed the phenotypes observed in nephrocytes overexpressing Piezo wildtype and might therefore be a potential intervention to prevent Piezo associated phenotypes.

## Discussion

Mechanotransducers play a crucial role in podocyte biology and within this study, we describe the role of the mechanosensitive ion channel Piezo in Drosophila nephrocytes **(Figure 6E)**. Here, we could show expression of Piezo1 in murine podocytes by immunofluorescence, localizing within foot processes and the cell body. This confirms RNAscope data showing Piezo1 expression in podocytes from a recently published study ^32^. The authors also report an upregulation of Piezo1 in their experimental hypertensive nephropathy model in podocytes ^32^, illustrating the need to investigate both, loss and overexpression of the channel, to characterize its functional role. Another recent publication could also show expression of Piezo1 in podocytes and an increase of Piezo1 in podocytes in lupus nephritis ^33^. Our data shows that both, depletion and overexpression of wildtype Piezo result in a severe pericardial nephrocyte phenotype. This includes morphological alterations of the nephrocyte diaphragm, which results in a less dense expression pattern of Duf (dNEPH) and Pyd (dZO1), and functional disturbances, as assessed by FITC-Albumin uptake assays. To understand the functional role of the Piezo protein in nephrocytes in greater detail, we also investigated a mutant fly strain lacking the mechanosensitive channel activity. Interestingly, this mutant does not present with a pericardial nephrocyte phenotype. Even after increasing expression levels further, morphology or function are not changed, suggesting that the observed phenotype is at least partially mediated via the mechanosensitive domain.

In addition to depletion and overexpression, we aimed to unravel the effects of activating endogenous Piezo in the presence of mechanical force. To do so, we used Yoda1 to pharmacologically activate endogenous Piezo. Here, we could show a filtration defect after Yoda1 treatment and applying a mechanical stimulus. This confirms our hypothesis of Piezo expression in nephrocytes and its importance in nephrocyte biology. We further confirmed this hypothesis by performing all Yoda1 experiments in a Piezo knockdown background, which abrogates the observed filtration defect.

Within this study, we also performed Ca^++^ imaging in isolated pericardial cells from adult flies and could show an increase of intracellular Ca^++^ levels after applying mechanical stimuli and pretreatment with Yoda1. This elevated Ca^++^ influx was abrogated by depleting Piezo in nephrocytes, confirming that the influx is at least partially mediated via Piezo. Our data suggests an opening of the channel upon a mechanical stimulus (puff) in nephrocytes, which is facilitated after binding of Yoda1 between the Arm domain and the repeat A ^36^. In a recent study, Sivakumar et al. showed that nephrocytes exhibit spontaneous calcium signals *in vivo* ^37^, which was also observed here. Analysing peak amplitude parameters enabled us to distinguish between the spontaneous calcium signals and the signals elicited after applying a puff. Sivakumar et al. were also able to show a functional role of store operated Ca^++^ entry (SOCE) mechanisms, in particular Stim and Orai-dependent, on nephrocyte endocytosis ^37^. Thus, this data, together with the data provided in here, show that Ca^++^ signalling is an important signalling feature in *Drosophila* nephrocytes.

Of note, in about half of the flies in our study (treated either with DMSO or Yoda1), Ca^++^ levels did not return to baseline levels within the timeframe of the recordings. This data suggests, that the mechanical stimulus applied, modulates several different Ca^++^ transport (import and export) mechanisms (e.g. SOCE), which slow down the decrease of intracellular Ca^++^ levels.

Mechanosensors, such as TRP channels, have also been described to regulate GTPase activation in podocytes ^40^. One downstream target of GTPases and Ca^++^ signalling is the actin cytoskeleton, which plays an essential role during podocyte injury and rearrangement processes ^41^. A previous study described actin clusters and cortical actin in nephrocytes as well ^42^. Here, we also assessed the actin cytoskeleton after Yoda1 treatment and applying mechanical force as well as upon overexpression of Piezo wildtype. While activation of endogenous Piezo does not result in alterations of the actin cytoskeleton, overexpression of Piezo caused an increase of cortical actin fibres. In *in vitro* podocytes activation of Piezo also caused actin cytoskeleton remodelling as a result of Rac1 activation ^33^. Of note, the absence of a phenotype upon Yoda1 treatment and applying mechanical force in our experiment, might be explained by the time frame of the experiment. Longer exposure of Yoda1, for example by feeding, is very interesting, but beyond the scope of this study. However, the more artificial condition of overexpression of Piezo, suggests an impact on the actin cytoskeleton, which might be mediated by alterations in Ca^++^ influx, in nephrocytes.

Our findings, that actin fibre formation might have relevance to nephrocytes response to injury, are supported by a previous study investigating the effect of Bis-T-23 in podocytes. Bis-T-23 promotes actin-dependent dynamin oligomerization and stabilizes the actin cytoskeleton during podocyte injury ^43^. In detail, several injury models which caused actin-cytoskeleton rearrangement could be at least partially rescued by Bis-T-23 treatment, again emphasising the importance of the actin-cytoskeleton and upstream regulators such as GTPases ^43^.

Analysing publicly available data from the Kidney Precision Medicine Project (KPMP) revealed an increase of Piezo1 in podocytes in CKD samples, which is in line with data obtained from experimental hypertensive nephropathy and lupus nephritis mouse models ^32,33^. Hence, a treatment to block Piezo activation could be beneficial for patients. Our data showed that the treatment with tarantula toxin (GsMTx4) was able to, reverse the observed phenotypes in nephrocytes overexpressing Piezo wildtype. This is in line with data showing improved kidney function in a lupus mouse model after GsMTx4 treatment ^33^. In addition, another study investigated the association between Piezo1 activation and kidney fibrosis ^44^. Tubular Piezo1 was increased in fibrotic kidneys and inhibition of Piezo1 with GsMTx4 resulted in attenuated tubulointerstitial fibrosis ^44^, further supporting the idea of GsMTx4 use as a potential intervention in the future. However, we want to mention that GsMTx4 is not specific to Piezo, but also inhibits other mechanosensitive channels. Interestingly, Piezo2 has recently been described to be expressed in renin-producing cells of the juxta-glomerular apparatus and mesangial cells in the kidney, where it contributes to renal blood volume sensing ^45,46^.

Taken together, here we provide evidence that the mechanosensitive ion channel Piezo is expressed in podocytes and nephrocytes and that Piezo activity levels need to be tightly regulated in order to maintain normal morphology and function in adult pericardial nephrocytes.

## Methods

### Fly husbandry

The UAS-Gal4 system is influenced by temperature, being more active at a higher temperature, thus we used two different temperatures (medium: 25°C and high: 28°C) for our experiments. The nephrocyte-specific knockdown or overexpression was achieved by mating UAS fly strains with the *sns*-Gal4 strain. Fly strains are listed in **table 1**.

**Table 1:** List of fly strains.

| **Fly strain** | **Origin** | **Purpose** | **Chromosome** |
| --- | --- | --- | --- |
| UAS-*piezo*-flag | BDSC ID: 95296 | Overexpression | 3. Chr. |
| UAS-*piezo*.2306MYC.flag | BDSC ID: 95298 | Overexpression | 3. Chr. |
| UAS-*piezo*-RNAi | VDRC ID: 101915 | Knockdown | 2. Chr. |
| *sns*-Gal4; UAS-*dicer2* | the *sns*-Gal4 strain was obtained from S. Abmayr<sup>47</sup> and combined with UAS-*dicer2*<sup>8</sup> | Nephrocyte-specific driver | 2. + 3. Chr. |
| UAS-GCaMP6medium | BDSC ID: 42750 | Overexpression | 3. Chr. |
| W<sup>1118</sup> |  | wildtype |  |

### Yoda1 and GsMTx4 treatment

To activate Piezo, nephrocytes were incubated with the specific activator compound Yoda1 for 5mins (30μM in DMSO; control: DMSO) (Sigma, now Merck). A mechanical stimulus was applied using the pressure injector SYS-PV830 (W.P.I. Instruments, Friedberg, Germany) in combination with a 1ml syringe fitted with a 26G needle as applicator nozzle. The syringe was either filled with Yoda1 (30µM) or DMSO (equal amount to Yoda1) and positioned with the same angle and position relative to the bath solution’s surface throughout the experiments. The tissue was incubated in the respective bath solution for 5mins prior to applying the mechanical stimulus. This was achieved by pressure-controlled release of the syringe solution (referred to as a puff, 0.2bar for 3secs) to simulate the induction of shear stress. To inhibit mechanosensitive channel activity, nephrocytes were incubated with the unspecific cationic mechanosensitive channel inhibitor tarantula toxin/GsMTx4 for 5mins (5μM in H_2_O; control: HL3.1 buffer) (Alomone labs, Jerusalem, Israel, Cat. nr: STG-100).

### Immunofluorescences of *Drosophila* tissue

Garland nephrocytes were isolated from 3^rd^ instar larvae in HL3.1 buffer. For isolation of pericardial cells, the cuticle of the abdomen of 1-3 days old flies were dissected leaving pericardial cells still attached to the heart tube. The whole tissue was used for immunofluorescence staining and the dissection was performed in HL3.1 buffer. After isolation, cells were fixed in 4% formaldehyde for 20mins and one hour in methanol. Primary and secondary antibody (see **table 2**) incubation was done according to standard procedures overnight at 4°C. Imaging was done using either a Zeiss LSM 800 confocal combined with an Airyscan for higher resolution or a Leica SP8 confocal. Images were further processed using ImageJ (version 1.53c). To quantify nephrocyte density (nephrocyte diaphragm length [μm/μm^2^]) we used our previously published macro for FIJI, which was adapted for Airyscan confocal images ^48^.

**Table 2:** List of antibodies.

| **Name** | **Company/ Provider** | **Catalog nr. / Reference** | **Host Species** | **Dilution IF** |
| --- | --- | --- | --- | --- |
| Duf | M. Ruiz-Gomez | <sup>28</sup> | Rabbit | 1:100 |
| Pyd | Developmental Studies Hybridoma Bank | PYD2 | Mouse | 1:25 |
| Piezo1 | Proteintech | 28511-1-AP | Rabbit | 1:100 |
| Synaptopodin | Synaptic Systems | 163 004 | Guinea Pig | 1:200 |

### Immunofluorescence of murine tissue

Paraffin-embedded tissue from wildtype mice was used for immunofluorescence staining. Tissue was de-paraffinized by passing through a descending alcohol row. After washing for three times (5 mins) with H2O followed by three washing steps with Dako wash buffer (Agilent Technologies, Santa Clara, USA), antigen retrieval was done in a steamer using pH9 antigen retrieval from Dako. Primary antibodies (listed in Table 1) were diluted in Dako antibody diluent and incubated overnight at 4°C. The next day sections were washed for three times (5 mins) with Dako wash buffer, followed by secondary antibody incubation with DAPI for one hour at room temperature. Prior to mounting with Prolong Gold, samples were again washed three times. Imaging of samples was done at a Zeiss LSM 900 confocal combined with Airyscan. Images were further processed using ImageJ (version 1.53c).

### FITC-Albumin uptake assay

FITC-Albumin uptake assays were performed as previously published ^8,49^. Uptake of FITC-Albumin into the lacunae is a read-out for nephrocyte function. Pericardial cells were isolated from 1-3 days old adults. Imaging was done with a Zeiss LSM 5 confocal, a Leica SP8 confocal and a Zeiss LSM 800 confocal microscope and images were further processed and analysed using ImageJ (version 1.53c). For comparative analysis, the exposure time of experiments was kept identical. The mean intensity of FITC-Albumin was measured and normalized to control cells.

### Visualization and quantification of actin fibres

To visualize actin fibres, we used phalloidin. For actin quantification cortical and surface mean intensities of phalloidin were measured. Images were obtained using a Zeiss LSM800 confocal microscope combined with Airyscan and further image processing was done using ImageJ (version 1.53c). For comparative analysis, laser intensity and settings were kept equal. Quantification of cortical fibres was done using the brush selection tool from FIJI.

### Ca^++^ imaging of pericardial nephrocytes

For Ca^++^ imaging experiments, UAS-GCaMP6m flies were combined with the nephrocyte-driver *sns*-Gal4 to obtain experimental animals. Flies were kept at 25°C until the experimental flies were 1-3 days old. Flies were anesthetized with Flynap (Carolina Biological Supply company, North Carolina, USA). Pericardial nephrocytes were isolated but still attached to the heart tube and the cuticle. The tissue was glued onto coverslips with vaseline and transferred into a recording chamber. The experimental setup was consistent with the previously described experiments (Yoda1 treatment). The recording bath was filled with either 1ml Yoda1 (30µM) in HL3.1 buffer or an equivalent volume of DMSO in HL3.1 buffer. The tissue was incubated in the respective bath solution for 5mins before applying a pressure stimulus. Using the same method as before, a 1ml syringe with a 26G needle, filled with either Yoda1 (30µM) or DMSO, was positioned on the surface of the bath solution and the syringe solution was released as a puff (0.2bar for 3secs, PDES-01 a.m., NPI Electronic GmbH, Tamm, Germany) to simulate shear stress. GCaMP6 was excited at 488nm, and fluorescence was collected between 500 and 530nm using a confocal microscope (eC1, Nikon, Düsseldorf, Germany). Time series of images were recorded at a rate of 1.26 frame per second. Data was evaluated with Nikon EZ-C1 Viewer (Nikon), processed using Excel (Microsoft, USA), and statistical tests were applied with Graph Pad Prism software. GCaMP6m-expressing nephrocytes were marked as a region of interest (ROI) and the mean fluorescence intensity within the ROI was measured throughout the time series. The mean fluorescence intensity (F) was normalized to the baseline fluorescence intensity before the puff (mean of 10 values prior to puff) and set to 1.0, hence changes in fluorescence intensity are stated as ΔF. Quantification of Ca^++^ transients (peak amplitude) was achieved by calculating the maximum increase of GCaMP6 intensity after the puff within 25secs after puff. The number of experiments is given as n, where n represents the averaged values of all ROIs analysed in one fly and equals one datapoint in the graphs. The figure legends include a depiction of how many flies and ROIs per condition were investigated. Traces of calcium signals were created with OriginPro 9.1G (OriginLab Corporation, USA) and composed using Adobe Illustrator and Adobe Photoshop (Adobe Systems Software, Ireland). Visualizations of changes in fluorescence were created using Fiji ImageJ (National Institutes of Health, USA) and composed with Adobe Photoshop (Adobe Systems Software, Ireland). In these visualizations, red signals indicate the difference between baseline brightness and brightness at the indicated timeslots (ΔF), the grey background signal represents baseline fluorescence prior to applied stimuli.

### Transcriptomic data analysis

Publicly available human single nucleus RNA-sequencing (snRNAseq) ^34^ data were searched for Piezo1 expression. Data and analyser have been used provided by KIT (kidney interactive transcriptomics) http://humphreyslab.com/SingleCell/. Permission to use the graphs generated has been provided by Benjamin Humphreys. In addition, human single cell RNA-sequencing data (scRNAseq) provided by the Kidney Precision Medicine Project (KPMP) consortium were searched for Piezo1 expression. The results here are in whole or part based upon data generated by the Kidney Precision Medicine Project. Accessed November 14th, 2023. https://www.kpmp.org. Funded by the National Institute of Diabetes and Digestive and Kidney Diseases (Grant numbers: U01DK133081, U01DK133091, U01DK133092, U01DK133093, U01DK133095, U01DK133097, U01DK114866, U01DK114908, U01DK133090, U01DK133113, U01DK133766, U01DK133768, U01DK114907, U01DK114920, U01DK114923, U01DK114933, U24DK114886, UH3DK114926, UH3DK114861, UH3DK114915, UH3DK114937).

### Statistical analysis

To determine statistical significance Graph Pad Prism software version 8 for Mac (GraphPad Software, San Diego, CA) was used. All results are expressed as means ± SEM. Datapoints represent single flies from 2 to 3 independent matings. Mean values from all imaged nephrocytes of one fly have been calculated and are shown as one datapoint in the graphs. Comparison of two groups was done using a student’s t-test, while comparison of more than two groups with one independent variable was done using one-way ANOVA followed by Tukey’s multiple comparisons test. A *P-value* < 0.05 was considered statistically significant.

## Acknowledgement

The Anti-Pyd monoclonal antibody developed by Fanning ^50^ was obtained from the Developmental Studies Hybridoma Bank, created by the NICHD of the NIH and maintained at The University of Iowa, Department of Biology, Iowa City, IA 52242. Images were created with BioRender.com.

## Author contribution

S.K. conceived the study, made the figures and wrote the paper, K.S., P.H.-C., K.K.A., I.E. and S.K. performed experiments, S.K., M.L. and K.S. analysed data, B.D., C.L. and T.B.H. revised the study critically for important intellectual content, S.K. and B.D. revised the paper; all authors approved the final version of the manuscript.

## Data Availability Statement

Data sharing not applicable as no datasets were generated during this study.

